# Insights into LIS1 function in cargo-adapter-mediated dynein activation *in vivo*

**DOI:** 10.1101/683995

**Authors:** Rongde Qiu, Jun Zhang, Xin Xiang

## Abstract

Deficiency of the LIS1 protein causes lissencephaly, a brain developmental disorder. Although LIS1 binds the microtubule motor cytoplasmic dynein and has been linked to dynein function in many experimental systems, its mechanism of action remains unclear. Here we revealed the function of LIS1 in cargo-adapter-mediated dynein activation in the model organism *Aspergillus nidulans*. Specifically, we found that overexpressed cargo adapter HookA (Hook in *A. nidulans*) missing its cargo-binding domain (ΔC-HookA) causes dynein and its regulator dynactin to relocate from the microtubule plus ends to the minus ends, and this dramatic relocation requires LIS1 and its binding protein NudE. Astonishingly, the requirement for LIS1 or NudE can be bypassed to a significant extent by specific mutations that open the auto-inhibited “phi-dynein” in which the motor domains of the dynein dimer are held close together. Our results suggest a novel mechanism of LIS1 action: it promotes the switch of dynein from the auto-inhibited state to an open state to facilitate dynein activation.

**Summary:** This study reveals the role of Lissencephaly 1 (LIS1) in cargo-adapter-mediated dynein activation. Furthermore, it discovers a novel mechanism of LIS1 action involving a switch of dynein from an auto-inhibited state to an active state.

## Introduction

Cytoplasmic dynein-1 (called “dynein” hereafter) is a microtubule motor that transports a variety of cargos in eukaryotic cells, and defects in dynein-mediated transport is linked to devastating neurodegenerative diseases and brain developmental disorders (Maday et al., 2014; Jaarsma and Hoogenraad, 2015; Bertipaglia et al., 2018). The dynein-cargo interaction requires the multi-component dynactin complex as well as specific cargo adapters (Schroer, 2004; Akhmanova and Hammer, 2010; Fu and Holzbaur, 2014; Reck-Peterson et al., 2018; Olenick and Holzbaur, 2019). Importantly, dynactin and cargo adapters also activate the motility of cytoplasmic dynein *in vitro* (McKenney et al., 2014; Schlager et al., 2014; Reck-Peterson et al., 2018; Olenick and Holzbaur, 2019). The mechanism underlying this activation was suggested by a recent cyro-EM analysis (Zhang et al., 2017a). Specifically, the two motor domains of the dynein heavy chain (HC) dimer are held together in an inactive “phi” conformation (Torisawa et al., 2014; Zhang et al., 2017a), which is in equilibrium with an “open” conformation in which the two domains are separated, and the binding to dynactin and cargo adapter causes the HC dimer to become parallel for directional movement along microtubules (Zhang et al., 2017a). Moreover, some cargo adapters facilitate the recruitment of a second dynein dimer to dynactin (Grotjahn et al., 2018; Urnavicius et al., 2018), further enhancing dynein force and speed (Urnavicius et al., 2018). While these are important steps toward understanding dynein regulatory mechanisms, the cargo-adapter-mediated dynein activation has never been analyzed *in vivo*, and it is especially unclear whether this process is regulated by other proteins *in vivo*.

Two of the most well-known yet enigmatic dynein regulators are LIS1 and its binding partner NudE (Kardon and Vale, 2009; Vallee et al., 2012; Reck-Peterson et al., 2018; Olenick and Holzbaur, 2019). LIS1 is encoded by the *lis1* gene whose deficiency causes type I lissencephaly, a human brain developmental disorder (Reiner et al., 1993). Fungal genetic studies first linked LIS1 to dynein function (Xiang et al., 1995a; Geiser et al., 1997; Willins et al., 1997). In the filamentous fungus *Aspergillus nidulans*, the LIS1 homolog NudF is critical for dynein-mediated nuclear distribution (Xiang et al., 1995a). *A. nidulans* genetics also led to the identification of NudE, a NudF/LIS1-binding protein (Efimov and Morris, 2000; Feng et al., 2000; Niethammer et al., 2000; Sasaki et al., 2000), whose homologs participate in dynein function in various systems (Minke et al., 1999; Liang et al., 2004; Li et al., 2005; Liang et al., 2007; Stehman et al., 2007; Yamada et al., 2008; Kardon and Vale, 2009; Ma et al., 2009; Zhang et al., 2009; Pandey and Smith, 2011; Wang and Zheng, 2011; Zylkiewicz et al., 2011; Vallee et al., 2012; Wang et al., 2013; Klinman and Holzbaur, 2015; Kuijpers et al., 2016; Olenick and Holzbaur, 2019). The mechanisms of these two proteins have been studied intensively but mechanistically how they regulate dynein still remains unclear. The dynein HC contains six AAA domains in its motor ring (King, 2000; Asai and Koonce, 2001), and LIS1 binds AAA4 and regions near AAA4 (Huang et al., 2012; Toropova et al., 2014; DeSantis et al., 2017). Intriguingly, purified LIS1 inhibits dynein motility *in vitro* (Yamada et al., 2008; McKenney et al., 2010; Huang et al., 2012), unless ATP hydrolysis at AAA3 is blocked (DeSantis et al., 2017). NudE/Nudel (also called NDEL1) relieves the inhibitory effect of LIS1 (Yamada et al., 2008; Torisawa et al., 2011), and the NudE-LIS1 complex enhances dynein force production (McKenney et al., 2010; Reddy et al., 2016). Moreover, dynactin partially relieves the inhibition of LIS1 on dynein motility (Wang et al., 2013), and when both dynactin and cargo adapter are present, LIS1 no longer inhibits but mildly enhances dynein movement (Baumbach et al., 2017; Gutierrez et al., 2017; Jha et al., 2017). All these studies have clearly demonstrated that the function of LIS1 is context-dependent. However, the exact molecular mechanism of LIS1 action on dynein regulation is not known (Reck-Peterson et al., 2018; Olenick and Holzbaur, 2019).

We have been using the fungal model organism *A. nidulans* to investigate dynein regulation *in vivo*. Unlike budding yeast where dynein is required almost exclusively for nuclear migration/spindle orientation (Eshel et al., 1993; Li et al., 1993), dynein, dynactin and LIS1 in filamentous fungi are required not only for nuclear distribution (Plamann et al., 1994; Xiang et al., 1994; Xiang et al., 1995a) but also for transporting a variety of other cargos including early endosomes and their hitchhiking partners (Wedlich-Soldner et al., 2002; Lenz et al., 2006; Abenza et al., 2009; Zekert and Fischer, 2009; Baumann et al., 2012; Bielska et al., 2014a; Higuchi et al., 2014; Egan et al., 2015; Guimaraes et al., 2015; Pohlmann et al., 2015; Salogiannis et al., 2016; Penalva et al., 2017; Otamendi et al., 2019). In filamentous fungi and budding yeast, dynein, dynactin and LIS1-NudE all accumulate at the microtubule (MT) plus ends (Han et al., 2001; Efimov, 2003; Lee et al., 2003; Sheeman et al., 2003; Zhang et al., 2003; Li et al., 2005; Lenz et al., 2006; Moore et al., 2008; Callejas-Negrete et al., 2015). The MT plus-end accumulation of dynein is important for spindle-orientation/nuclear migration and for early endosome transport (Lee et al., 2003; Sheeman et al., 2003; Lenz et al., 2006; Omer et al., 2018; Xiang, 2018). In *A. nidulans* and *Ustilago maydis*, the plus-end dynein accumulation depends on dynactin and kinesin-1 but not NudF/LIS1 (Zhang et al., 2003; Lenz et al., 2006; Zhang et al., 2010; Egan et al., 2012; Yao et al., 2012). This differs from the situation in budding yeast where LIS1 is critical for dynein’s plus-end accumulation and in mammalian cells where both LIS1 and dynactin are critical (Lee et al., 2003; Sheeman et al., 2003; Splinter et al., 2012). Recently, the interaction between fungal dynein and early endosome has been found to be mediated by dynactin as well as the Fhip-Hook-Fts (FHF) complex (Walenta et al., 2001; Xu et al., 2008; Zhang et al., 2011; Bielska et al., 2014b; Zhang et al., 2014). Within the FHF complex, Hook (HookA in *A. nidulans* and Hok1 in *U. maydis*) interacts with dynein-dynactin and Fhip interacts with early endosome (Bielska et al., 2014b; Yao et al., 2014; Zhang et al., 2014; Guo et al., 2016; Schroeder and Vale, 2016). The function of dynactin and the Hook complex in early endosome transport is evolutionarily conserved (although multiple Hook proteins in mammalian cells participate in even more functions of dynein) (Yeh et al., 2012; Guo et al., 2016; Dwivedi et al., 2019; Olenick et al., 2019), and importantly, mammalian Hook proteins activate dynein *in vitro* (McKenney et al., 2014; Olenick et al., 2016; Schroeder and Vale, 2016).

Here we developed a new assay to examine HookA-mediated dynein activation in *A. nidulans* and revealed the role of LIS1 in this context. Specifically, we overexpressed HookA lacking the C-terminal early endosome-binding site (ΔC-HookA), which binds dynein-dynactin but not early endosome (Zhang et al., 2014). In contrast to the MT plus-end accumulation of dynein-dynactin in wild-type cells, ΔC-HookA overexpression shifts the accumulation to the MT minus ends. LIS1 and its binding protein NudE are both required for this cargo-adapter-mediated dynein relocation *in vivo*. Interestingly, dynein mutations that open up the inactive phi conformation of dynein bypass the requirement of LIS1 or NudE to a significant extent *in vivo*. We discuss these results to suggest a novel link between the function of LIS1 to a key step of dynein activation: shifting from an auto-inhibited phi-conformation to an open conformation that allows dynein to be fully activated.

## Results

### Dynein and dynactin are relocated from the MT plus ends to the minus ends upon overexpression of ΔC-HookA

To examine cargo-adapter-mediated dynein activation *in vivo*, we sought to create *A. nidulans* cells where dynein and dynactin are occupied by cytosolic cargo adapters. To do that, we replaced the wild-type *hookA* allele with the *gpdA*-ΔC-*hookA*-S allele so that ΔC-HookA (missing its cargo-binding site) is overexpressed under the constitutive *gpdA* promoter (Figure 1A, B) (Pantazopoulou and Penalva, 2009; Zhang et al., 2011). Overexpression of ΔC-HookA did not inhibit colony growth significantly (Figure 1C) but caused a partial nuclear-distribution defect (sFigure 1). In wild-type strains, GFP-labelled dynein and dynactin (p150 subunit) formed comet-like structures near the hyphal tip, representing their MT plus-end accumulation (Han et al., 2001; Zhang et al., 2003) (Figure 1D, E, F). Upon overexpression of ΔC-HookA, the plus-end comets of dynein or dynactin disappeared (Figure 1D, E, F). This was not caused by a defect in MT organization because mCherry-labelled ClipA/Clip170 (Zeng et al., 2014) formed plus-end comets in the same cells (Figure 1D). A dominant feature in these cells is the accumulation of dynein and dynactin at septa, which are structures known to contain active microtubule-organizing centers (MTOCs) (Konzack et al., 2005; Xiong and Oakley, 2009; Zekert et al., 2010; Zhang et al., 2017b) (Figure 1E, F). Dynein at septa was found in wild-type cells (Liu et al., 2003), but the signals were much less obvious compared to the strong accumulation upon ΔC-HookA overexpression (Figure 1E). Besides septal MTOCs, the nuclear envelope-associated spindle-pole body (SPB) represents the earliest-discovered MTOC (Oakley et al., 1990). To determine if dynein or dynactin is at the SPBs, we used the NLS-DsRed fusion that labels nuclei during interphase (Shen and Osmani, 2013). We found clear SPB-like signals of dynein and dynactin on some nuclei (Figure 1G, H), which were never observed in wild-type cells. However, the septal signals of dynein-dynactin in the *gpdA*-ΔC-*hookA*-S cells were brighter and more consistently observed than the SPB signals, and thus, we used the septal signals to indicate MT minus-end accumulation in the rest of the work.

**Figure 1.**
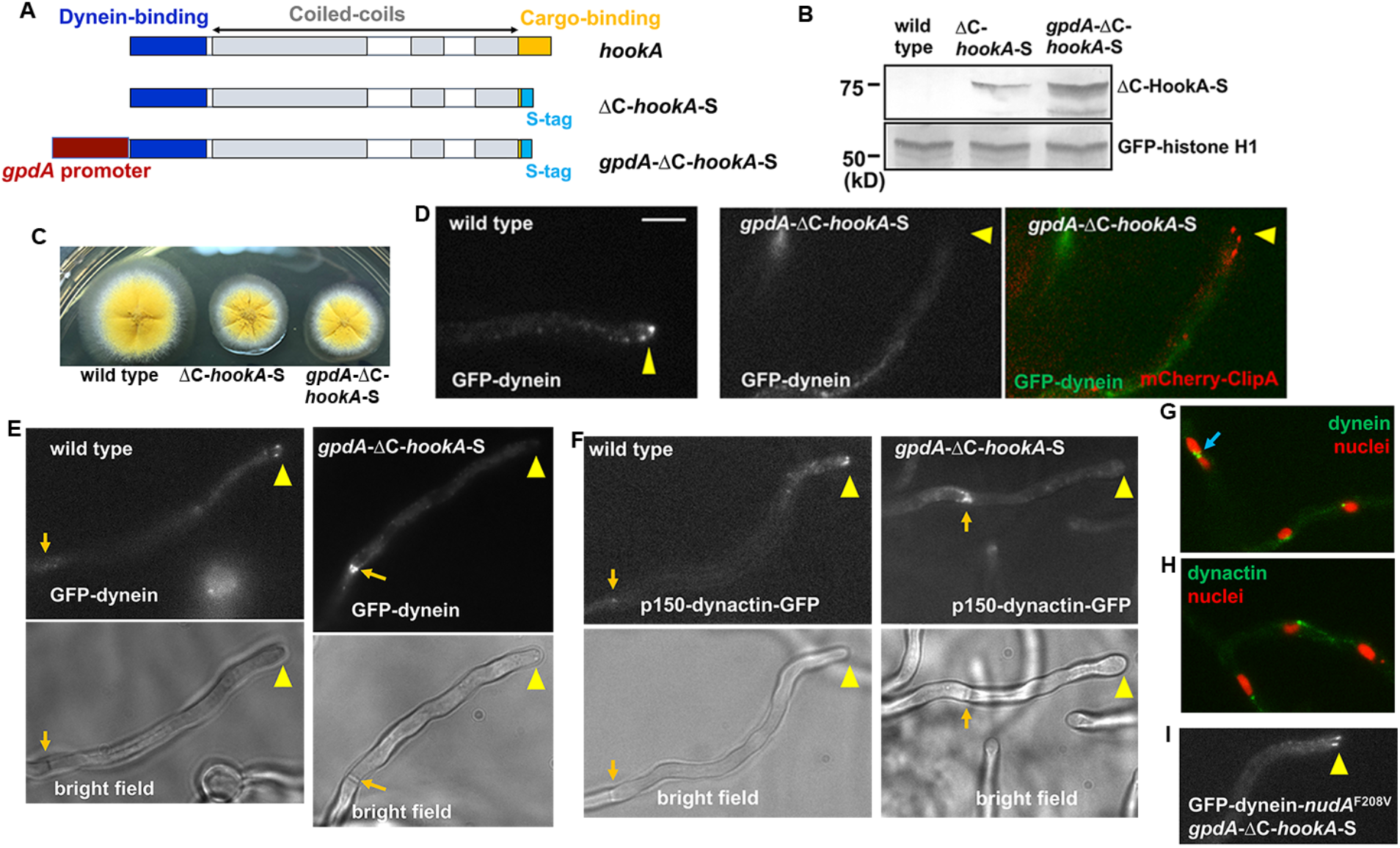
Overexpression of ΔC-HookA-S by the *gpdA* promoter drives dynein-dynactin relocation from the MT plus ends to the minus ends. (A) A diagram showing the wild-type *hookA* allele, the ΔC-*hookA*-S allele and the *gpdA*-ΔC-*hookA*-S allele. (B) Western blots showing that the ΔC-HookA-S protein is overexpressed by the *gpdA* promoter. GFP-Histone H1 was used as a loading control. (C) The colony phenotype of *gpdA*-ΔC-*hookA*-S strain in comparison to that of a wild-type control and a strain containing the ΔC-*hookA*-S allele. (D) Images of GFP-dynein HC in wild type and *gpdA*-ΔC-*hookA*-S cells. Plus-end dynein comets near the hyphal tip (yellow arrowhead) are present in wild type but not in the *gpdA*-ΔC-*hookA*-S strain, but the plus-end comets of mCherry-ClipA are present in the same *gpdA*-ΔC-*hookA*-S cell. Bar, 5 μm. (E) Accumulation of GFP-dynein on septum (light brown arrow) in the *gpdA*-ΔC-*hookA*-S strain. Bright-field images are shown below to indicate hyphal shape and position of septum. (F) Septal accumulation of dynactin p150-GFP in the *gpdA*-ΔC-*hookA*-S strain. (G) GFP-dynein localization on nuclei labelled with NLS-DsRed in the *gpdA*-ΔC-*hookA*-S strain. A blue arrow indicates dynein in the middle of two closely-associated nuclei. (H) Dynactin p150-GFP localization on nuclei labelled with NLS-DsRed in the *gpdA*-ΔC-*hookA*-S strain. (I) Plus-end dynein comets in the nudAF^208V^, *gpdA*-ΔC-*hookA*-S strain.

Although the plus-end to minus-end (MTOC) relocation of dynein-dynactin is fully consistent with cargo adapter-mediated dynein activation observed *in vitro* (McKenney et al., 2014; Schlager et al., 2014; Olenick et al., 2016; Schroeder and Vale, 2016), we sought to further confirm that this relocation needs functional dynein. To do that, we examined the effect of a previously identified dynein loss-of-function mutation, *nudA*^F208V^. The *nudA*^F208V^ mutation in the dynein tail impairs dynein-mediated nuclear distribution and early endosome transport but does not affect dynein-dynactin interaction or dynein-early endosome interaction (Qiu et al., 2013), consistent with the importance of the tail in dynein motor activity (Ori-McKenney et al., 2010; Rao et al., 2013; Hoang et al., 2017). Upon overexpression of ΔC-HookA, plus-end comets formed by GFP-dynein with the *nudA*^F208V^ mutation were still detected (Figure 1I), confirming that functional dynein is needed for this relocation.

### NudF/LIS1 is required for ΔC-HookA-activated relocation of dynein from the MT plus ends to the minus ends

To examine the role of NudF/LIS1 in ΔC-HookA-mediated dynein activation *in vivo*, we used the temperature-sensitive (ts) *nudF6* mutant in which the NudF/LIS1 protein is unstable at its restrictive temperature (Xiang et al., 1995a). The *nudF6* mutation (identified as *nudF*^L304S^ in this work) caused a significant defect in ΔC-HookA-activated relocation of dynein and dynactin from the MT plus ends to the septa (Figure 2A, B, C). In sharp contrast to the disappearance of plus-end dynein or dynactin comets and the septal accumulation of these proteins in *gpdA-ΔC-* HookA-S cells, the bright plus-end comets of dynein-dynactin were easily observed in the *nudF6, gpdA*-ΔC-HookA-S cells, and no septal accumulation was observed (Figure 2A, B, C). Similar to *nudF6*, another ts mutation *nudF7* (Xiang et al., 1995a) and a conditional-null mutation *alcA-nudF* that allows NudF/LIS1 expression to be shut off by glucose-mediated repression at the *alcA* promoter (Xiang et al., 1995a) also retained dynein at the MT plus ends upon overexpression of ΔC-HookA (Figure 2D). Together, these results indicate that NudF/LIS1 is critically required for ΔC-HookA-mediated dynein activation in *A. nidulans*.

**Figure 2.**
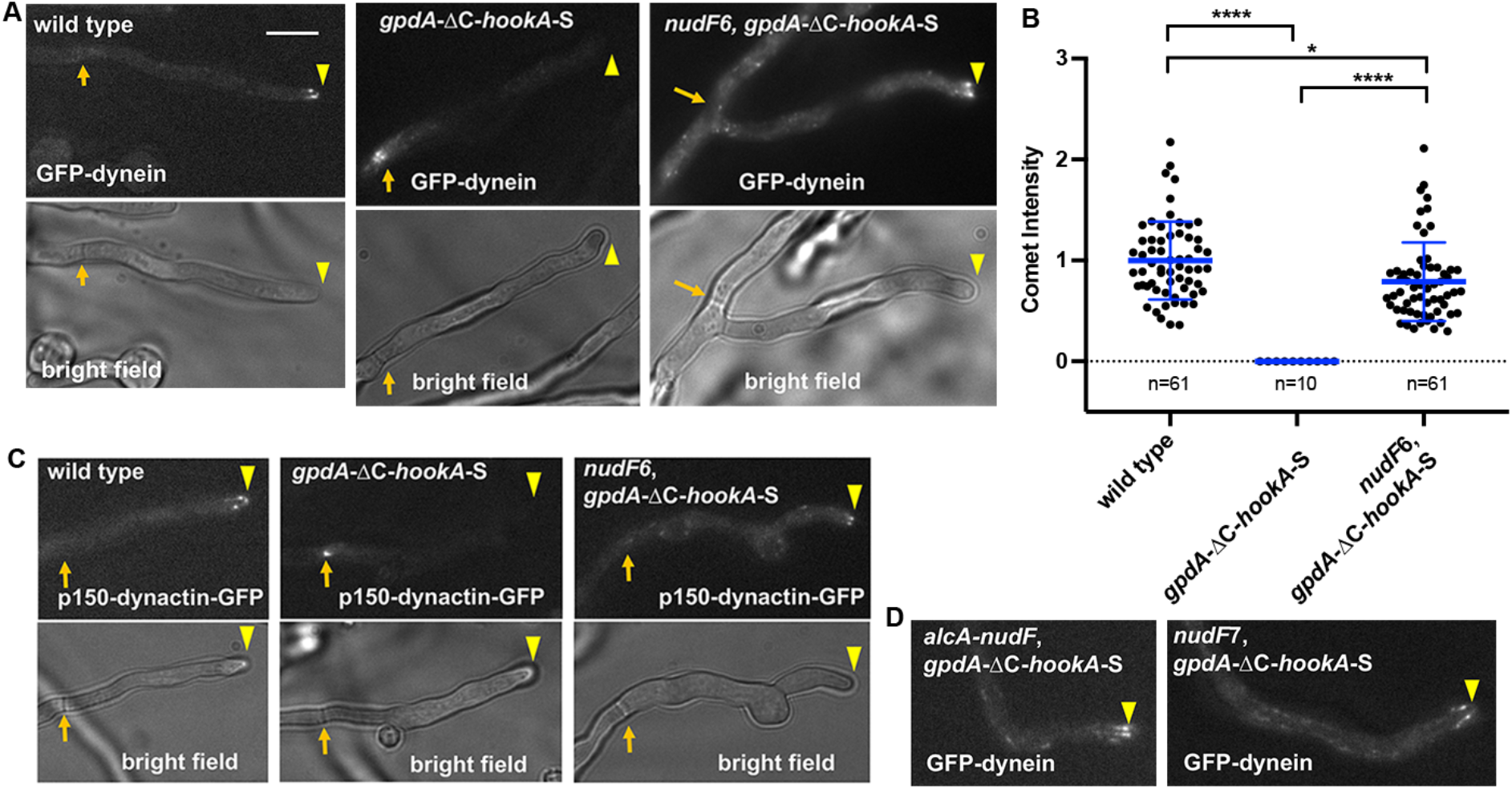
NudF/LIS1 is required for the relocation of dynein and dynactin from the MT plus ends to the minus ends upon ΔC-HookA overexpression. (A) GFP-dynein in wild type, a *gpdA*-ΔC-*hookA*-S strain and a *nudF6, gpdA*-ΔC-*hookA*-S strain. Note that GFP-dynein accumulates as MT plus-end comets but not at septum in the *nudF6, gpdA*-ΔC-*hookA*-S strain. Bright-field images are shown below to indicate hyphal shape and position of septum. Hyphal tip is indicated by a yellow arrowhead and septum by a light brown arrow. Bar, 5 μm. (B) A quantitative analysis on dynein comet intensity in the *nudF6*, gpdA-ΔC-HookA-S strain in comparison to those in wild-type and the gpdA-ΔC-HookA-S strain. The average wild-type value is set as 1. Scatter plots with mean and S.D. values were generated by using Prism 7. * indicates p<0.05 and **** indicates p<0.0001 (one-way ANOVA, unpaired). (C) Dynactin p150-GFP in wild type, a *gpdA*-ΔC-*hookA*-S strain and a *nudF6, gpdA*-ΔC-*hookA*-S strain. Note that GFP-dynactin accumulates as plus-end comets but not at septum in the *nudF6, gpdA*-ΔC-*hookA*-S strain. Bright-field images are shown below to indicate hyphal shape and position of septum. Hyphal tip is indicated by the yellow arrowhead and septum by a light brown arrow. (D) GFP-dynein in a *nudF7, gpdA*-ΔC-*hookA*-S strain and an *alcA-nudF, gpdA*-ΔC-*hookA*-S strain. Note that GFP-dynein plus-end comets are present in these cells.

### NudF/LIS1 does not accumulate at the MT minus ends upon overexpression of ΔC-HookA

Unlike dynein or dynactin, NudF/LIS1-GFP did not accumulate at septa upon overexpression of ΔC-HookA (Figure 3A). Instead, the plus-end comets formed by NudF/LIS1-GFP were still observed upon overexpression of ΔC-HookA, although the intensity was significantly lower than that in wild-type cells (Figure 3A, B). Thus, ΔC-HookA may drive some NudF/LIS1 proteins to leave the MT plus end with dynein-dynactin, but the association is not maintained. This is consistent with previous studies showing that dynein and dynactin associate with the early endosome undergoing dynein-mediated transport but LIS1 dissociates from it after the initiation of transport from the MT plus ends (Lenz et al., 2006; Egan et al., 2012).

**Figure 3.**
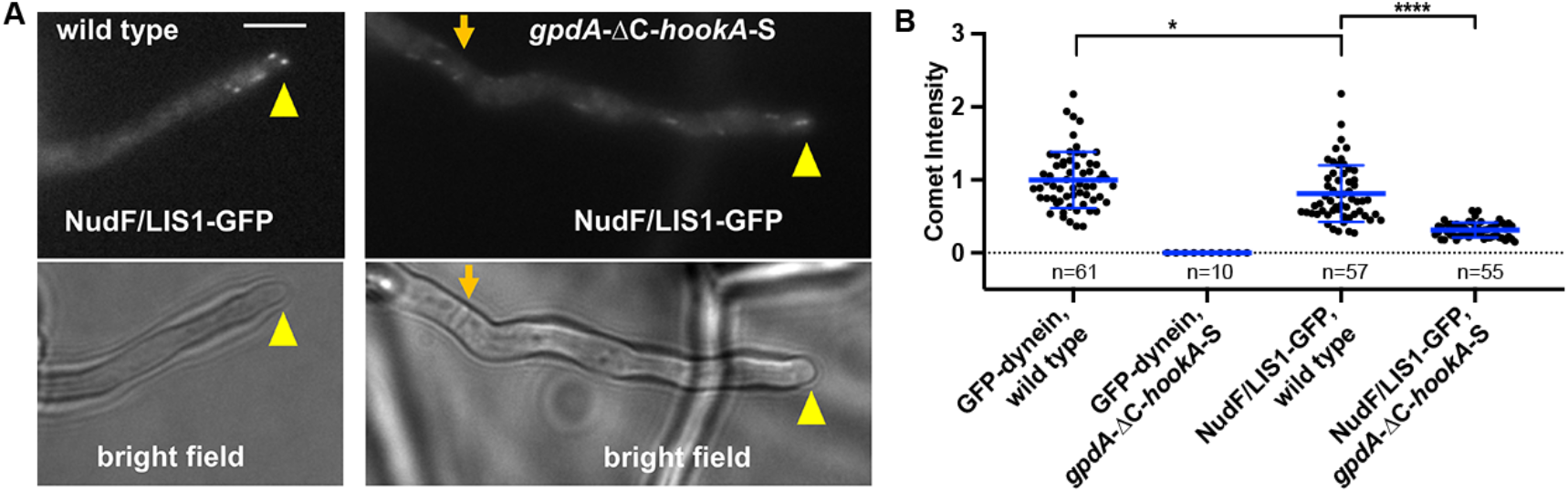
NudF/LIS1 proteins do not relocate to the MT minus ends upon overexpression of ΔC-HookA. (A) NudF/LIS1-GFP signals are seen as plus-end comets and not accumulated on septa in cells containing *gpdA*-ΔC-*hookA*-S. Hyphal tip is indicated by a yellow arrowhead and septum by a light brown arrow. Bar, 5 μm. (B) A quantitative analysis of the NudF/LIS1-GFP comet intensity in a *gpdA*-ΔC-*hookA*-S strain in comparison to that in the wild type. In this analysis, we also added the data for GFP-dynein as used in Figure 2B for comparison. The average wild-type value for GFP-dynein is set as 1. Scatter plots with mean and S.D. values were generated by using Prism 7. * indicates p<0.05 and **** indicates p<0.0001 (one-way ANOVA, unpaired).

### NudF/LIS1 is important for dynein activation not only at the MT plus ends

We next sought to address whether dynein can undergo cargo adapter-mediated activation only at the MT plus ends. In fungi and higher eukaryotic cells including neurons, plus-end-directed kinesins are required for the accumulation of dynein at the MT plus ends (Zhang et al., 2003; Carvalho et al., 2004; Lenz et al., 2006; Arimoto et al., 2011; Roberts et al., 2014; Twelvetrees et al., 2016). In the Δ*kinA* (Kinesin-1) mutant of *A. nidulans* (Requena et al., 2001), dynein fails to arrive at the MT plus ends but locates along MTs (Zhang et al., 2003; Zhang et al., 2010; Egan et al., 2012). We found that GFP-dynein in cells with the Δ*kinA* and *gpdA*-ΔC-*hookA*-S alleles accumulated at septa (Figure 4A), suggesting that dynein along MTs can be activated before arriving at the plus ends as long as cargo adapters are available globally. However, adding the *nudF6* mutation to the genetic background with the Δ*kinA* and *gpdA*-ΔC-*hookA*-S alleles caused the septal accumulation of dynein to be significantly decreased and dynein along MTs to be more obvious (Figure 4A, B). Thus, NudF/LIS1 is important for cargo-adapter-mediated dynein activation even when dynein fails to arrive at the MT plus ends. These results could be relevant to the previous findings that LIS1 is required for dynein-mediated cargo transport in both the distal and mid-axon of cultured neuronal cells and that minus-end-directed cargo transport can start in the middle of a microtubule if the cargo has the opportunity to meet and bind dynein along the MT in *U. maydis* (Schuster et al., 2011; Moughamian et al., 2013).

**Figure 4.**
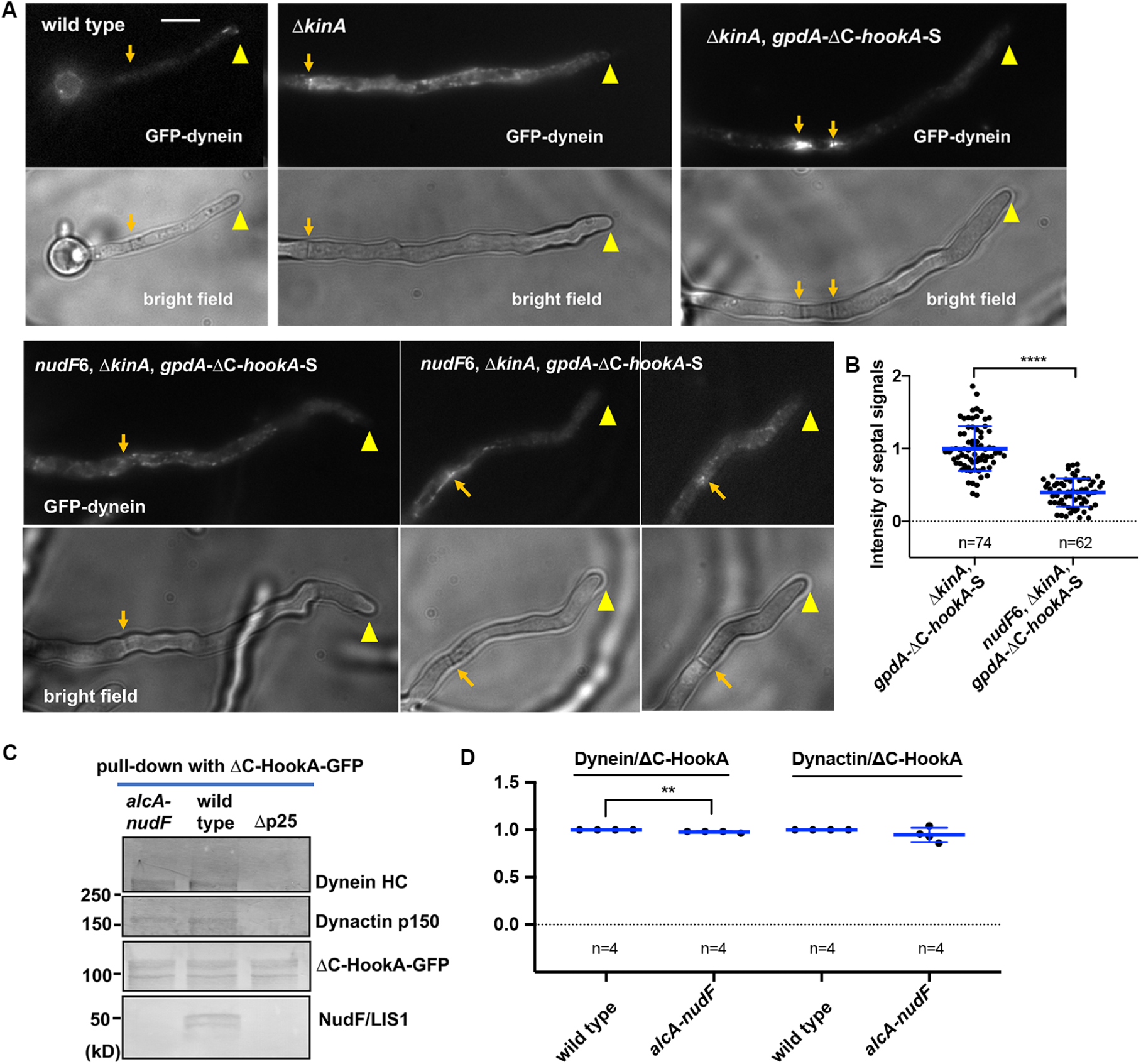
Dynein along MTs in the Δ*kinA* mutant relocates to the minus ends upon ΔC-HookA overexpression, and NudF/LIS1 plays an important role in this process. (A) Images of GFP-dynein in wild type, a Δ*kinA* strain, a Δ*kinA, gpdA-ΔC-hookA-S* strain and a *nudF6, ΔkinA, gpdA-ΔC-hookA-S* strain. Bright-field images are shown below to indicate hyphal shape and position of septum. Hyphal tip is indicated by a yellow arrowhead and septum by a light brown arrow. (B) A quantitative analysis of the septal GFP-dynein signal intensity. The average value for the Δ*kinA, gpdA-ΔC-hookA-S* strain is set as 1. Scatter plots with mean and S.D. values were generated by using Prism 7. **** indicates p<0.0001 (student *t*-test, unpaired). (C) Western blots showing that dynein HC and dynactin p150 are pulled down with ΔC-HookA-GFP in wild type and the *alcA-nudF* mutant in which the expression of NudF/LIS1 is shut off. (D) A quantitative analysis on the ratio of pulled-down dynein HC to ΔC-HookA-GFP (dynein/ΔC-HookA) and that of pulled-down dynactin p150 to ΔC-HookA-GFP (dynactin/ΔC-HookA). The values were generated from western analyses of four independent pull-down experiments. The wild-type values are set as 1. Scatter plots with mean and S.D. values were generated by using Prism 7. ** indicates p<0.01 (student *t*-test, unpaired).

### The dynein-dynactin-ΔC-HookA complex is still formed without NudF/LIS1

We next used a biochemical pull-down assay to determine if the defect in dynein activation in cells lacking NudF/LIS1 is caused by a defect in the formation of the dynein-dynactin-ΔC-HookA complex. For this assay, we combined the ΔC-HookA-GFP fusion (Zhang et al., 2014) with the *alcA-nudF* mutant where NudF/LIS1 expression is shut off by glucose-mediated repression at the *alcA* promoter (Xiang et al., 1995a). The Δp25 mutant was used as a negative control because p25 of dynactin is important for the formation of the dynein-dynactin-ΔC-HookA complex (Zhang et al., 2014; Qiu et al., 2018). In the *alcA-nudF* mutant where NudF/LIS1 is undetectable, both dynactin and dynein were still pulled down with ΔC-HookA-GFP, and only a very mild reduction in the ratio of pulled-down dynein to ΔC-HookA compared to the wild type was detected (Figure 4C, D). Thus, the role of NudF/LIS1 in dynein activation cannot be simply explained by its involvement in the formation of the dynein-dynactin-cargo adapter complex *in vivo*, suggesting that NudF/LIS1 must be critical for another step of dynein activation.

### The phi-opening mutations allow the requirement of NudF/LIS1 to be bypassed to a significant extent

A key step of dynein activation is the switch of dynein from the auto-inhibited phi-dynein conformation (Torisawa et al., 2014; Zhang et al., 2017a) to an open conformation that can readily undergo activation by dynactin and cargo adapter *in vitro* (Zhang et al., 2017a). Formation of phi-dynein depends on ionic interactions between the dynein HC linker domain and AAA4, and mutating two linker residues opens up phi-dynein and causes dynein to accumulate at centrosomes/spindle poles in mammalian cells (Zhang et al., 2017a). To investigate the relationship between phi-dynein and LIS1 function, we constructed an *A. nidulans* strain containing the two analogous phi-opening mutations *nudA*^R1602E, K1645E^ (Figure 5A). In addition, we also obtained the *nudA*^R1602E^ mutant containing only one of the two phi-opening mutations due to a homologous recombination between the two sites. In either case, the wild-type *nudA* allele is replaced by the mutant allele (confirmed by sequencing of the genomic DNA). While the *nudA*^R1602E^ mutant formed a colony that appeared normal, the *nudA*^R1602E, K1645E^ mutant formed a smaller colony especially at a higher temperature such as 37°C or 42°C (Figure 5B, sFigure 2), and it also showed a partial defect in nuclear distribution (sFigure 2).

**Figure 5.**
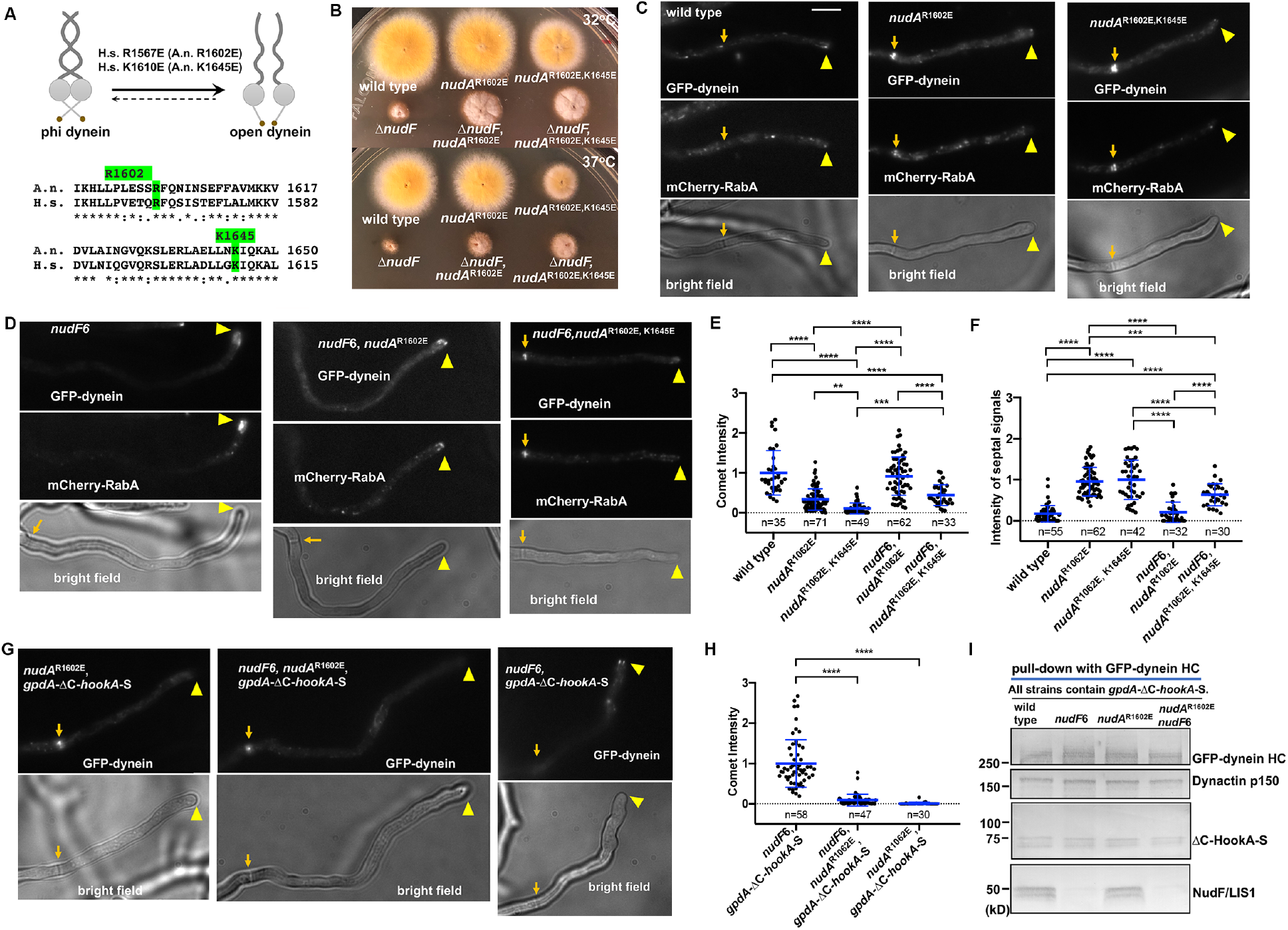
Phi-opening partially bypasses NudF/LIS1 function. (A) A cartoon showing mutations that open the phi dynein (Zhang et al., 2017a) and a sequence alignment showing the corresponding dynein HC regions from human (H. s.) and *A. nidulans* (A. n.). (B) Colony phenotypes of phi-opening mutants and suppression of the Δ*nudF* growth defect by the phi-opening mutations at 32°C and 37°C. (C) GFP-dynein with *nudA^R1602E^* or *nudA*^R1602E, K1645E^ accumulates at septa with mCherry-RabA-labelled early endosomes. Hyphal tip is indicated by a yellow arrowhead and septum by a light brown arrow. Bar, 5 μm. (D) Images of GFP-dynein and mCherry-RabA in a *nudF6* strain, a *nudF6, nudA*^R1602E^ strain and a *nudF6, nudA*^R1602E, K1645E^ strain. Note that GFP-dynein with *nudA*^R1602E, K1645E^ accumulates at septa with mCherry-RabA-labelled early endosomes in the *nudF6* mutant. (E) A quantitative analysis on GFP-dynein comet intensity. The average value for the wild-type strain is set as 1. (F) A quantitative analysis on the septal GFP-dynein signal intensity. The average value for the *nudA*^R1602E, K1645E^ strain is set as 1. For both (E) and (F), scatter plots with mean and S.D. values were generated by using Prism 7. **** indicates p<0.0001, *** indicates p<0.001, ** indicates p<0.01 (one-way ANOVA, unpaired). (G) The *nudA*^R1602E^ mutation causes GFP-dynein HC to accumulate at the septa in *gpdA*-ΔC-*hookA*-S cells regardless of NudF/LIS1 function. (H) A quantitative analysis on dynein comet intensity in strains shown in (G). **** indicates p<0.0001. (I) Western blots showing that the *nudA*^R1602E^ mutation does not apparently affect the amounts of dynactin p150 and ΔC-HookA-S pulled down with GFP-dynein.

The *nudA*^R1602E, K1645E^ mutant exhibited an striking accumulation of GFP-dynein at septa together with mCherry-RabA-labeled early endosomes (Figure 5C). Introducing the Δ*hookA* allele into this background caused the plus-end dynein comets to reappear but abolished the colocalization of early endosomes with dynein at septa (sFigure 3) (note that the septal dynein did not completely disappear, suggesting the existence of other cargo adapters in *A. nidulans* that could also activate dynein) (sFigure 3). Thus, GFP-dynein with *nudA*^R1602E, K1645E^ is able to arrive at the MT plus end where it undergoes robust activation to relocate to the minus end. Interestingly, GFP-dynein with the *nudA*^R1602E^ single mutation also accumulated at septa together with early endosomes (Figure 5C), suggesting that this mutation must have opened the phi-dynein at least partially to cause a more robust dynein activation.

To determine if the phi-opening mutations can bypass NudF/LIS1 function *in vivo*, we introduced these mutations into the Δ*nudF* (nudF-deletion) mutant background. Amazingly, both the *nudA*^R1602E^ and *nudA*^R1602E, K1645E^ mutations enhanced growth of the Δ*nudF* mutant colony (Figure 5B). Moreover, the *nudA*^R1602E, K1645E^ mutations allowed dynein and early endosomes to be seen at septa (sFigure 3), suggesting that artificially opening the phi-dynein allows the requirement of NudF/LIS1 to be partially bypassed. We then performed a more detailed imaging analysis using the ts *nudF6* mutant containing either *nudA*^R1602E^ or *nudA*^R1602E, K1645E^ (note that the *nudF6* mutant is much healthier than Δ*nudF* at a lower temperature, allowing us to obtain enough spores for quantitative imaging). In the *nudF6* mutant grown at its restrictive temperature, GFP-dynein with *nudA*^R1602E^ mainly formed plus-end comets, but GFP-dynein with *nudA*^R1602E, K1645E^ accumulated at septa together with early endosomes even though plus-end comets did not completely disappear (Figure 5D, E, F). This result suggests that the *nudA*^R1602E^ single mutation must have only opened phi dynein partially. However, upon overexpression of ΔC-HookA, GFP-dynein with *nudA*^R1602^ accumulated at septa in the *nudF6* background, which is in sharp contrast to the plus-end accumulation of wild-type dynein in the same genetic background (Figure 5G, H). This result suggests that even a partial opening of the phi conformation compensates for NudF/LIS1 function during cargo-adapter-mediated dynein activation. Interestingly, although phi-opening enhanced the formation of the mammalian dynein-dynactin-cargo adapter complex *in vitro* (Zhang et al., 2017a), the *nudA*^R1602E^ single mutation did not show any obvious enhancement of the dynein-dynactin-ΔC-HookA-S complex formation in *A. nidulans* (Figure 5I). Thus, we interpret these results to suggest that phi-opening *per se* is a key function of NudF/LIS1.

### NudE is required for dynein activation and this requirement is partially bypassed by the phi-opening mutations

Previous studies have suggested a role of NudE in enhancing LIS1 function by recruiting LIS1 to dynein (Efimov, 2003; Shu et al., 2004; Li et al., 2005; McKenney et al., 2010; Zylkiewicz et al., 2011; Wang et al., 2013). However, NudE was also thought to relieve the inhibition of LIS1 on dynein motility (Yamada et al., 2008). Moreover, the function of NudE in cargo-adapter-mediated dynein activation has never been addressed *in vivo* or *in vitro*. To address this, we first introduced GFP-dynein and the *gpdA*-ΔC-*hookA*-S allele into the Δ*nudE* background. We found that although overexpression of ΔC-HookA drives dynein relocation from the MT plus ends to the minus ends, this does not happen in the Δ*nudE* mutant (Figure 6A). Thus, just like NudF/LIS1, NudE is also required for cargo-adapter-mediated dynein activation.

**Figure 6.**
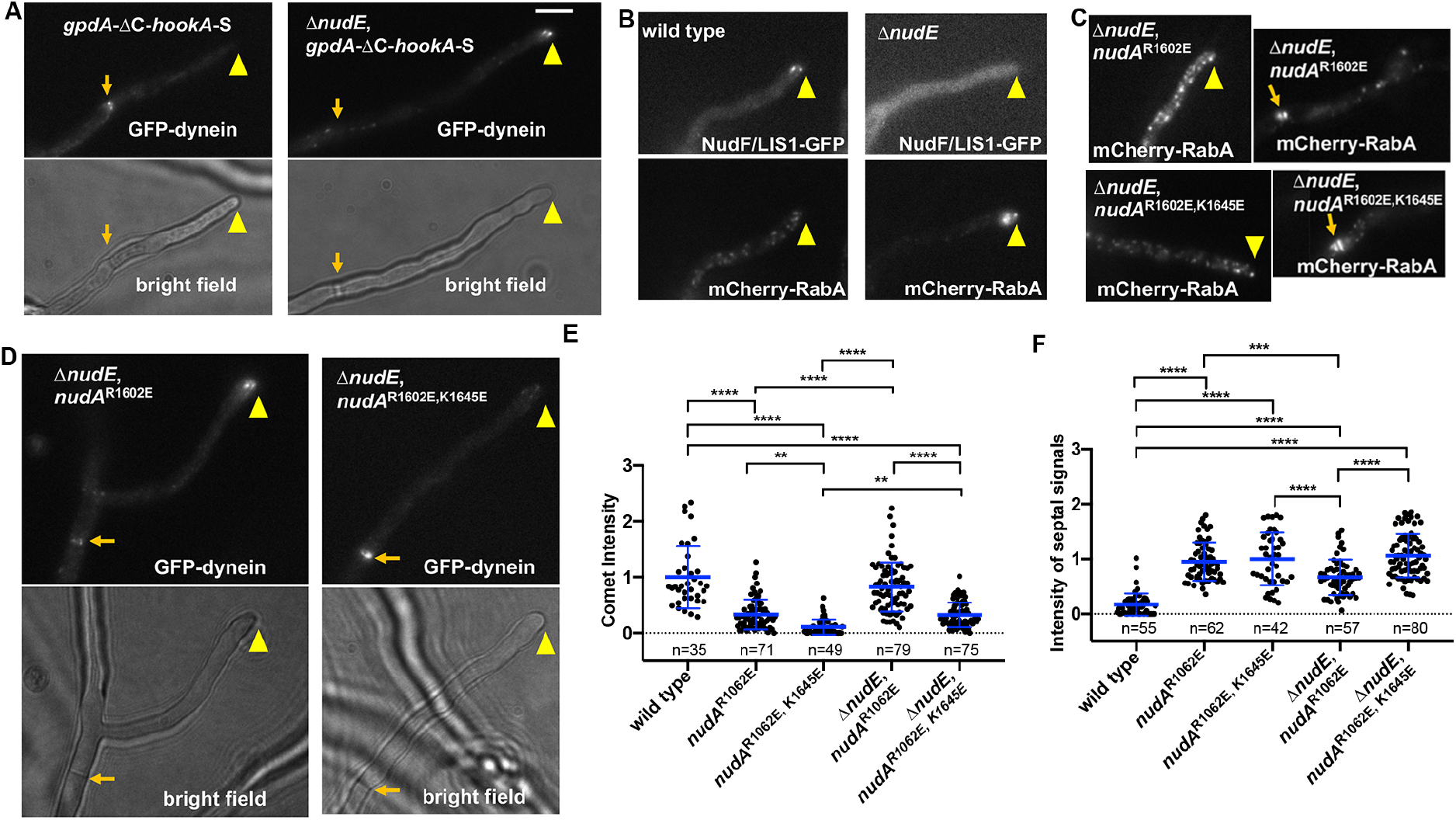
NudE is required for dynein activation and its function is bypassed by the phi-opening mutations. (A) GFP-dynein in a *gpdA*-ΔC-*hookA*-S strain and a Δ*nudE, gpdA-ΔC-hookA-S* strain. Hyphal tip is indicated by a yellow arrowhead and septum by a light brown arrow. Bar, 5 μm. (B) NudF/LIS1-GFP and mCherry-RabA in wild type and a Δ*nudE* strain. (C) mCherry-RabA in a Δ*nudE*, *nudA*^R1602E^ strain and a Δ*nudE, nudA*^R1602E, K1645E^ strain. (D) GFP-dynein in a Δ*nudE, nudA*^R1602E^ strain and a Δ*nudE, nudA*^R1602E, K1645E^ strain. Bright-field images are shown below to indicate hyphal shape and position of septum. (E) A quantitative analysis on dynein comet intensity. The average value for the wild-type strain is set as 1. (F) A quantitative analysis on septal GFP-dynein signal intensity. The average value for the *nudA*^R1602E, K1645E^ strain is set as 1. For both (E) and (F), scatter plots with mean and S.D. values were generated by using Prism 7. **** indicates p<0.0001, *** indicates p<0.001, ** indicates p<0.01 (one-way ANOVA, unpaired).

Furthermore, by observing the newly constructed NudF/LIS1-GFP fusion under its endogenous promoter, we revealed a clear role of NudE in NudF/LIS1 localization. Specifically, the plus-end NudF/LIS1-GFP comets in wild-type cells were no longer observed in the Δ*nudE* mutant and the GFP signals appeared diffused (Figure 6B). Previously, this role of NudE in NudF/LIS1 localization was only observed upon loss of ClipA/Clip170 when the experiment was done using the alcA-promoter-driven GFP-NudF/LIS1 fusion, possibly because the Δ*nudE* phenotype was less severe when NudF/LIS1 is expressed at a level higher than normal (Efimov, 2003; Efimov et al., 2006). We next addressed whether NudE is required for activating dynein that failed to arrive at the plus end, using the Δ*kinA* (kinesin-1) mutant. We found that while GFP-dynein in Δ*kinA, gpdA*-ΔC-*hookA*-S cells accumulated at septa, introducing the Δ*nudE* allele into this genetic background caused a more obvious MT decoration of GFP-dynein, consistent with an activation defect (sFigure 4). Thus, NudE is required for cargo-adapter-mediated dynein activation at locations beyond the MT plus ends, just like NudF/LIS1 (Figure 4A, B).

The Δ*nudE* mutant showed an abnormal accumulation of mCherry-RabA-labelled early endosomes at the hyphal tips (Figure 6B), a defect similar to that caused by loss of NudF/LIS1 (Zhang et al., 2010; Egan et al., 2012). Importantly, the phi-opening mutations, especially *nudA*^R1602E, K1645E^, significantly reduced the number of hyphal tips containing this abnormal accumulation of early endosomes (from >90% in Δ*nudE* to <10% in the Δ*nudE*, *nudA*^R1602E, K1645E^ strain, n=50) (Figure 6C). In addition, the phi-opening mutations also caused an obvious septal accumulation of early endosomes (Figure 6C). GFP-dynein with either *nudA*^R1602E, K1645E^ or *nudA*^R1602E^ accumulated at the septa in the Δ*nudE* mutant, although plus-end comets did not completely disappear (Figure 6D, E, F). The result that the *nudA*^R1602E^ mutation allowed a significant septal accumulation of dynein in the Δ*nudE* mutant (Figure 6F) but not in the *nudF6* mutant (Figure 5F) is consistent with the notion that the function of NudF/LIS1 is not totally abolished upon loss of NudE (note that the Δ*nudE* mutant grows much better than the *nudF/lis1* mutants) (Efimov and Morris, 2000; Efimov et al., 2006). Together, these results support the notion that NudE supports the function of LIS1 in dynein activation, and its function can be bypassed to a significant extent by opening the phi-dynein.

## Discussion

In this study, we developed a robust *in vivo* assay for cargo-adapter-mediated dynein activation, which allowed us to dissect the function and mechanism of the dynein regulator NudF/LIS1 (called “LIS1” hereafter). We found that both LIS1 and its binding-protein NudE are critical for cargo-adapter-mediated dynein activation *in vivo*. Remarkably, the requirement for LIS1 or NudE *in vivo* is bypassed to a significant extent if the auto-inhibited phi-dynein is opened up artificially. Our results provide the first *in vivo* evidence to suggest that LIS1-NudE may promote the opening of phi-dynein, a key step of dynein activation.

LIS1 is required for cytoplasmic dynein function in many different cell types (Kardon and Vale, 2009; Vallee et al., 2012; Olenick and Holzbaur, 2019). Possibly, it also regulates dynein inside cilia and flagella where a phi-like state of dynein is present (Pedersen et al., 2007; Rompolas et al., 2012; Toropova et al., 2017; Roberts, 2018). Recent studies have shown that the function of LIS1 is context-dependent (Yamada et al., 2008; McKenney et al., 2010; Huang et al., 2012; Wang et al., 2013; Baumbach et al., 2017; DeSantis et al., 2017; Gutierrez et al., 2017; Jha et al., 2017). Purified LIS1 in the absence of dynactin and cargo adapter inhibits dynein motility *in vitro* (Yamada et al., 2008; McKenney et al., 2010; Huang et al., 2012). In budding yeast, LIS1 is required for the MT plus-end accumulation of dynein (Lee et al., 2003; Sheeman et al., 2003), and the inhibitory effect of LIS1 on dynein motility may help retain dynein at the plus end (Lammers and Markus, 2015). It is interesting to note that budding yeast dynein is active by itself (Reck-Peterson et al., 2006), which could be why LIS1’s inhibitory function is needed for keeping dynein at the plus end until cargo adapters become available (Lammers and Markus, 2015). In *A. nidulans* and *U. maydis*, dynein and dynactin still accumulate at the MT plus ends without LIS1 (Zhang et al., 2003; Lenz et al., 2006; Egan et al., 2012), possibly because dynein-dynactin arrived at the plus ends are not activated before interacting with cargo adapters. This has facilitated our dissection of LIS1’s role in the minus-end-directed motility of dynein. Our results on the positive role of LIS1 in dynein activation are consistent with the main conclusion obtained from recent *in vitro* motility assays (Baumbach et al., 2017; Gutierrez et al., 2017; Jha et al., 2017). Nevertheless, the *in vivo* results are more striking: While the dynein-dynactin-cargo adapter complex moves robustly towards the MT minus end in the absence of LIS1 *in vitro* (McKenney et al., 2014; Schlager et al., 2014), LIS1 is critical for the ΔC-HookA-activated dynein relocation *in vivo*. We postulate that the intracellular environment may require dynein to operate under higher tension and with more complicated regulations (note that tension applied to the dynein linker domain *in vitro* also alters the regulatory requirement for dynein motility (Nicholas et al., 2015)).

In *A. nidulans*, dynein carrying the phi-opening mutations dramatically accumulates at septal MTOCs with early endosomes, suggesting that opening the phi-dynein must have allowed dynein to either bind more robustly to dynactin and cargo adapter (Zhang et al., 2017a) or switch to the active conformation more effectively after cargo binding. Interestingly, only the two mutations that open the phi-dynein more completely can bypass the requirement of LIS1 function to allow dynein to be accumulated at the septa. However, the single *nudA*^R1602E^ mutation, which presumably opens phi incompletely, also bypasses the requirement of LIS1 when ΔC-HookA is overexpressed. Thus, cargo adapter binding can further promote the open state to compensate for LIS1 loss. This is consistent with the model that dynactin and cargo adapter further switch dynein to the active state, thereby preventing the equilibrium from being shifted towards the phi state (Zhang et al., 2017a). Without dynactin and cargo adapter, LIS1 may still promote the open state but dynein would not be fully functional to move along microtubules (Zhang et al., 2017a). Both phi-opening and LIS1 promote the dynein-dynactin-cargo adapter complex formation *in vitro* (Baumbach et al., 2017; Zhang et al., 2017a). However, this effect of LIS1 is not obvious in *A. nidulans* in the presence of the cytosolic ΔC-HookA, and the *nudA*^R1602E^ mutation compensated for the loss of LIS1 function in the context of ΔC-HookA overexpression without obviously enhancing the dynein-dynactin-cargo adapter complex formation. Together, these results suggest that phi-opening *per se* is one key function of LIS1 in *A. nidulans* (Figure 7).

**Figure 7.**
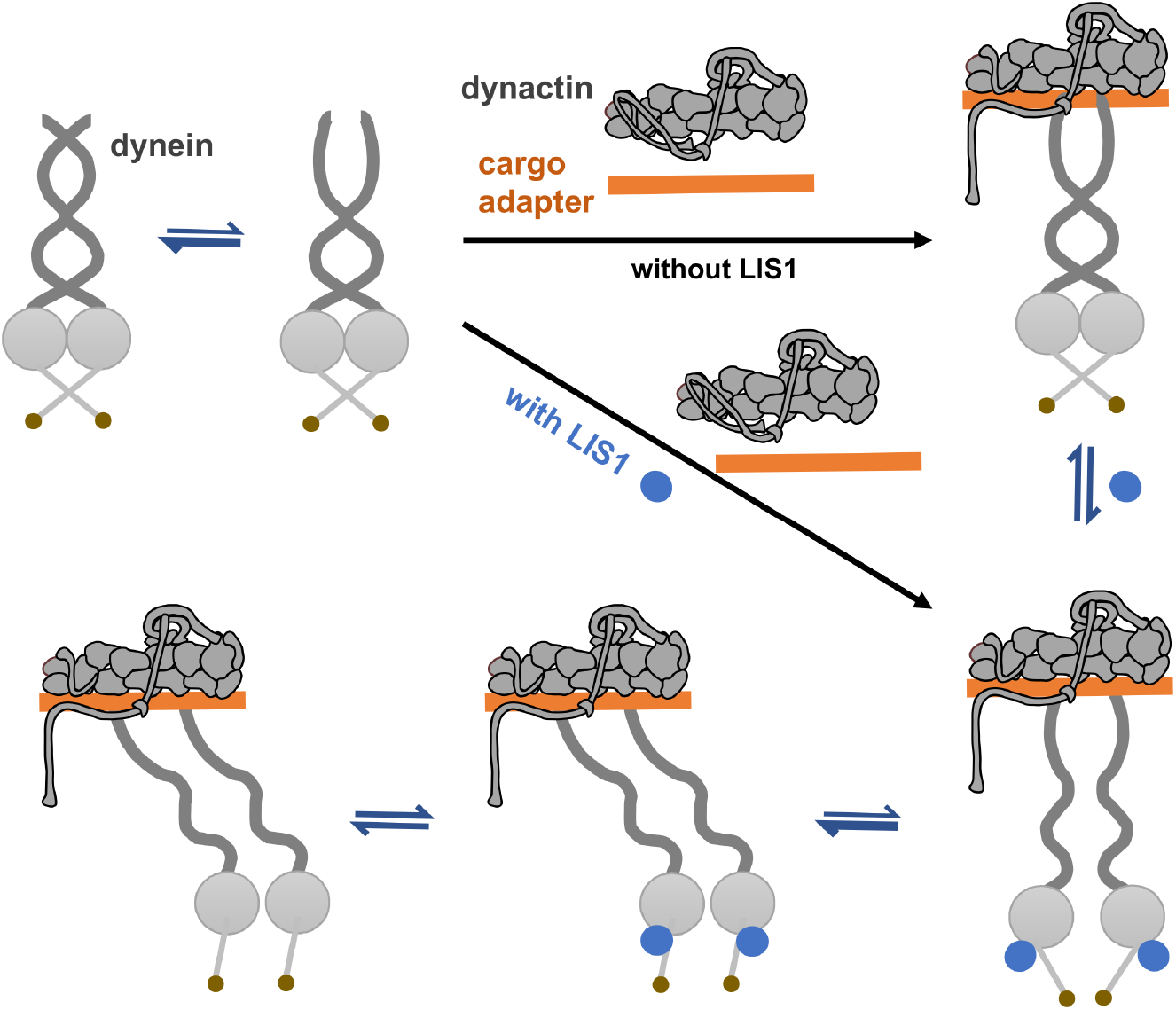
A model of LIS1 function in cargo-adapter-mediated dynein activation. Based on our result that the dynein-dynactin-ΔC-HookA complex is still formed in the absence of LIS1, we propose that between the fully closed and the fully open states of dynein, there is an intermediate state in which the dynein tails are partially open and able to bind dynactin and cargo adapters. However, this complex is not functional *in vivo* without LIS1. Based on our result that the phi-opening mutations mimic LIS1 function to a significant extent, we propose that LIS1 shifts the equilibria toward the fully open state and then toward the fully functional state where the two HCs within the dimer are parallel to each other (Zhang et al., 2017a). This fully functional state may have a relatively weak affinity for LIS1 to allow its dissociation from the complex at some point during the minus-end-directed movement.

Given the structural data that LIS1 binds to AAA4 and that the phi structure is maintained by ionic interactions between linker residues and AAA4 residues (Huang et al., 2012; Toropova et al., 2014; DeSantis et al., 2017; Zhang et al., 2017a), it is likely that LIS1 could bind the open dynein as an allosteric regulator to stabilize the open state (a similar hypothesis was also raised by other investigators independently (Olenick and Holzbaur, 2019)). This would in turn facilitate the cargo-adapter-mediated switch of dynein to a fully functional state with the two dynein HCs being in a parallel configuration (Zhang et al., 2017a). Interestingly, the LIS1-binding protein NudE is also required for dynein activation *in vivo*, and the requirement of NudE is also bypassed to a significant extent by the phi-opening mutations. This is consistent with the notion that NudE enhances LIS1 function. NudE binds to the dynein intermediate chain in the dynein tail, a site also required for dynactin binding (McKenney et al., 2011; Wang et al., 2013; Jie et al., 2017). As the two dynein tails are held together at several positions in the phi-dynein conformation (Figure 7) (Zhang et al., 2017a), it cannot be excluded that NudE may participate in phi-opening from the tail side to promote LIS1 function. However, because overexpression of LIS1 totally compensates for the loss of NudE (Efimov, 2003), we favor the possibility that NudE is not directly involved in phi-opening but simply helps bring LIS1 close to its site(s) of action.

Since the two phi-opening mutations allow LIS1 function to be bypassed to a significant extent but not completely, we would not rule out the possibility that LIS1 has additional roles in dynein regulation besides shifting the phi-dynein towards an open conformation. We should also point out that constitutively opening up the phi-dynein as achieved by the phi-opening mutations has a clear negative effect inside cells: It causes mitotic defects in mammalian cells (Zhang et al., 2017a), and it causes a defect in nuclear distribution in *A. nidulans* (sFigure 2). Thus, although our data point to the function of LIS1 in phi-opening, this process may be regulated by additional factors *in vivo*, including dynein’s own mechanical cycle, and identification of these regulatory factors will be an important task in the future.

## Materials and Methods

### Strains, media and live cell imaging

*A. nidulans* strains used in this study are listed in Table 1. Genetic crosses were done by standard methods, and progeny with desired genotypes were selected based on colony phenotype, imaging analysis, western analysis, diagnostic PCR and sequencing of specific regions of the genomic DNA. Media were described previously (Zhang et al., 2011). Fluorescence microscopy of live A. *nidulans* hyphae was as described (Zhang et al., 2014). All images were captured using an Olympus IX73 inverted fluorescence microscope linked to a PCO/Cooke Corporation Sensicam QE cooled CCD camera. An UplanApo 100x objective lens (oil) with a 1.35 numerical aperture was used. A filter wheel system with GFP/mCherry-ET Sputtered series with high transmission (Biovision Technologies) was used. The IPLab software was used for image acquisition and analysis. Quantitation of signal intensity was done as described previously (Zhang et al., 2014). Images were taken at room temperature immediately after the cells were taken out of the incubators. Cells were cultured overnight in minimal medium with 1% glycerol and supplements at 32°C or 37°C (all experiments using *nudF6* strains and controls were done at 37°C). Note that the *nudF6* mutant is temperature-sensitive (ts): it forms a tiny colony lacking asexual spores at a higher temperature (typical of a *nud* mutant), but some spores are produced at its semi-permissive temperature of 32°C. Thus, for experiments involving *nudF6*, we harvested spores at 32°C and cultured them at 37°C for imaging analysis. The *nudF6* mutant is much better than Δ*nudF* for imaging and biochemical analyses because we can harvest enough spores from the *nudF6* mutant at 32°C whereas the Δ*nudF* mutant is sick and does not produce spores at any temperature. For a few experiments using strains containing the *alcA-nudF* (conditional null) allele, we harvest spores from the solid minimal medium with 1% glycerol and cultured them in liquid minimal medium with 0.1% glucose for imaging analysis or YG (yeast extract and glucose) rich medium for protein pull-down experiments.

**Table 1.**
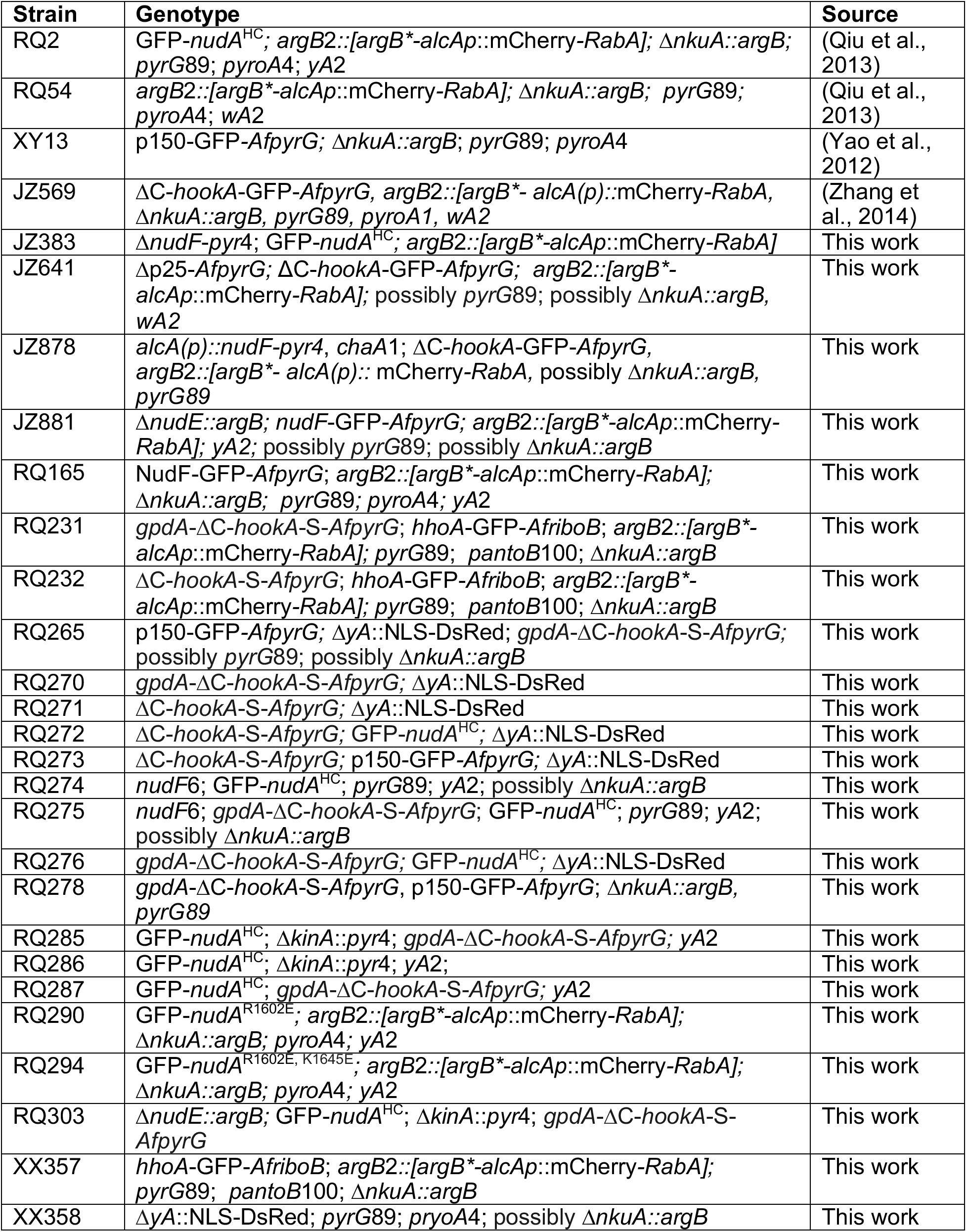
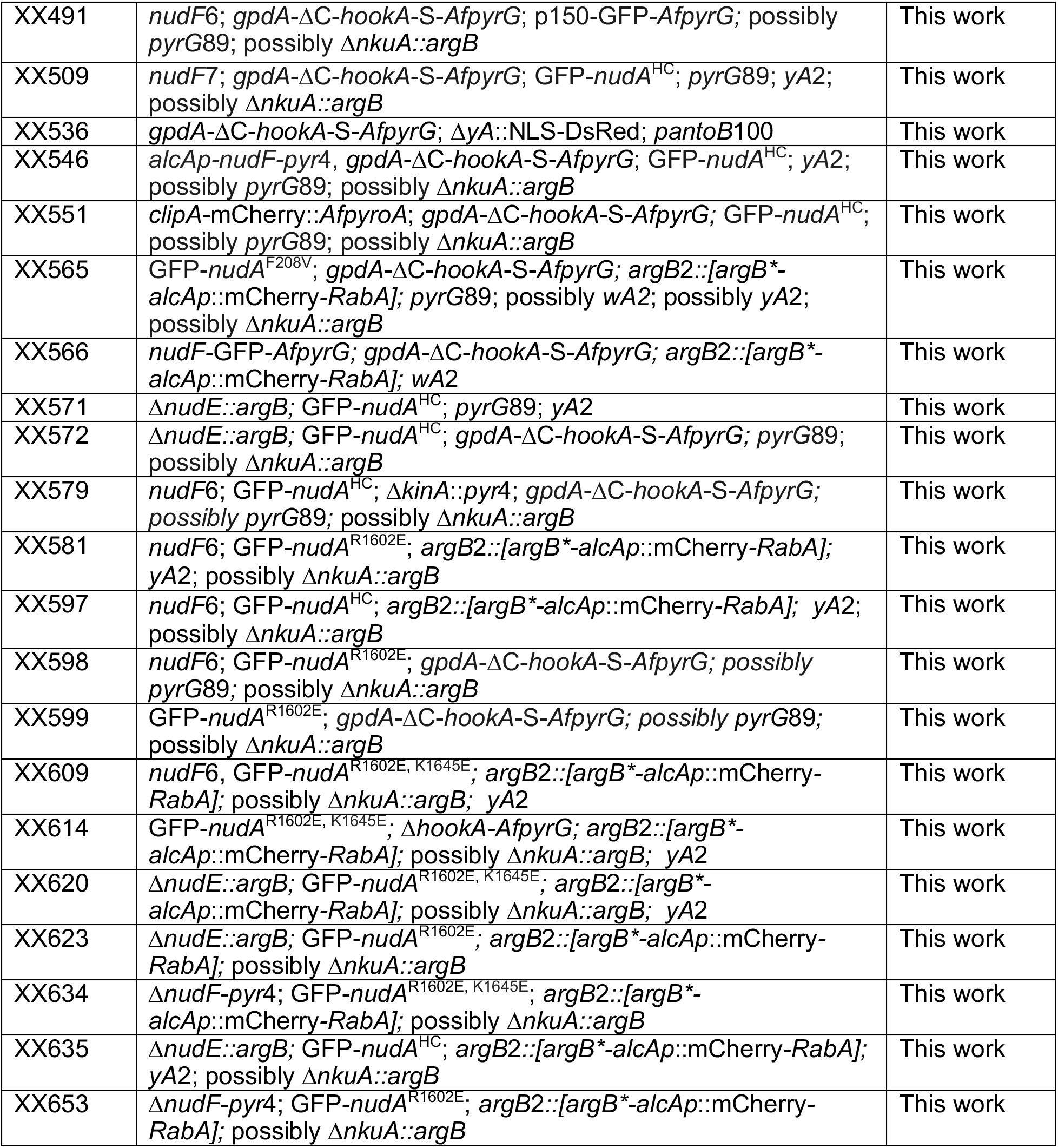
*Aspergillus nidulans* strains used in this study

### Constructing the *gpdA*-ΔC-*hookA*-S strain

For constructing the *gpdA*-ΔC-*hookA*-S strain, we co-transformed into the *A. nidulans* strain XX357 the fragment containing Δ*C-hookA-S-AfpyrG* (Qiu et al., 2018) with another fragment containing the ~1.2 kb *gpdA* promoter inserted in between the N-terminal HookA coding sequence and its upstream sequence. The *gpdA* promoter was inserted in this region using fusion PCR with primers 41U, HKpr2, gpdAF2, gpdAR2 (these four primers were described perviously(Zhang et al., 2014)), HKN2 (5’-AGTCGGAGCGTACCGT-3’) and HKgR (5’-TCAGCCTCAAGGTTTTGGTTC-3’).

### Constructing the *nudA*^R1602E^ and the *nudA*^R1602E, K1645E^ dynein HC “phi-opening” mutants

We used fusion PCR to make a DNA fragment of *nudA* containing both the R1602E and the K1645E mutations using the following primers: 1602F (5’-CGAGCGAGTT CCAGAAT AT CAACT CAGAATT CTT CG-3’), 1602R 5’-AGTTGATATTCTGGAACTCGCTCGATTCCAGGGGAAGAA-3’), 1645F (5’-GCT GCTT AACGAAAT CCAGAAAGCT CT CGGT GAAT AC-3’), 1645R (5’-GCTTTCTGGATTTCGTTAAGCAGCTCGGCCAG-3’), NudA54 (5’-GTGGATGAACTCATTCCAAGA-3’) and NudA36 5’-TTGGATCTACCAGCATAGCCA-3’). The fragment was co-transformed with a selective marker *pyrG* fragment into the RQ2 strain containing GFP-dynein HC (NudA) and mCherry-RabA. More than 200 *pyrG+* transformants were examined under the microscope, and several of them were selected because they exhibited clear septal enrichment of both the GFP and mCherry signals. Our sequencing analysis indicated that several strains contain both mutations (nudA^R1602Ei K1645E^), but one strain contains only the *nudA*^R1602E^ mutation due to homologous recombination between the two sites.

### Constructing the *nudF-GFP* strain

The following oligos, F5F (5’-ATCAGACTGGACGAAGCC-3’), F5R (5’-CAAATAGAATTAAATGGTACTCGAGTCGGTTGTTTGTGTTCGCAAAT-3’), gpdF (5’-GACTCGAGTACCATTTAATTCTATTTG-3’), gpdR (5’-TGTGATGTCTGCTCAAGCG-3), FF (5’-CGCTTGAGCAGACATCACAATGAGCCAAATATTGACAGCTCC-3’), FR (5’-GCCTGCACCAGCTCCGCTGAACACCCGTACAGAGTT-3’), GFPF (5’-GGAGCTGGTGCAGGC-3’), GFPR (5’-CTGTCTGAGAGGAGGCACTG-3’), F3F (5’-CAGTGCCTCCTCTCAGACAGGTCGCGATCTTCATCACAGTT-3’) and F3R (5’-CGACAGAATGGAACGGGAAA-3’) were used for fusion PCR to create the *gpdA-nudF-GFP-AfpyrG* fragment. This fragment was transformed into the RQ2 strain, and progeny with plus-end comets were selected. For this study, we only used a transformant (RQ165) containing NudF-GFP but not the gpdA-NudF-GFP fusion protein.

### Biochemical pull-down assays and western analysis

The μMACS GFP-tagged protein isolation kit (Miltenyi Biotec) was used to pull down proteins associated with the GFP-tagged protein. This was done as described previously (Zhang et al., 2014). About 0.4 g hyphal mass was harvested from overnight culture for each sample, and cell extracts were prepared using a lysis buffer containing 50 mM Tris-HCl, pH 8.0 and 10 μg/mL of a protease inhibitor cocktail (Sigma-Aldrich). Cell extracts were centrifuged at 8,000 *g* for 15 minutes and then 16,000 *g* for 15 minutes at 4°C, and supernatant was used for the pull-down experiment. To pull down GFP-tagged proteins, 25 μL anti-GFP MicroBeads were added into the cell extracts for each sample and incubated at 4°C for 30-40 minutes. The MicroBeads/cell extracts mixture was then applied to the μColumn followed by gentle wash with the lysis buffer used above for protein extraction (Miltenyi Biotec). Pre-heated (95°C) SDS-PAGE sample buffer was used as elution buffer. Western analyses were performed using the alkaline phosphatase system and blots were developed using the AP color development reagents from Bio-Rad. Quantitation of the protein band intensity was done using the IPLab software as described previously (Qiu et al., 2013). The antibody against GFP (polyclonal) was from Takara Bio Inc. The antibody against the S-tag was from Cell Signaling Technology. Affinity-purified antibodies against dynein HC, dynactin p150 and NudF/LIS1 were generated in previous studies (Xiang et al., 1995a; Xiang et al., 1995b; Zhang et al., 2008).

## Acknowledgements

We thank Reinhard Fischer, Bo Liu, Berl Oakley, Aysha Osmani, Stephen Osmani, Martin Egan, Samara Reck-Peterson, Miguel Peñalva and Xuanli Yao for sharing Aspergillus strains. We thank Stephen Osmani and Tian Jin for critical comments on the manuscript, and we thank Ahmet Yildiz and Samara Reck-Peterson for sharing unpublished results on LIS1. This work was funded by the National Institutes of Health grant R01GM121850-01A1 (to X.X.). The authors declare no competing financial interests.

## Author contributions

R. Qiu, J. Zhang and X. Xiang designed the experiments, performed the experiments and analyzed the data. X. Xiang wrote the paper with edits from R. Qiu and J. Zhang.

---

**sFigure 1.**
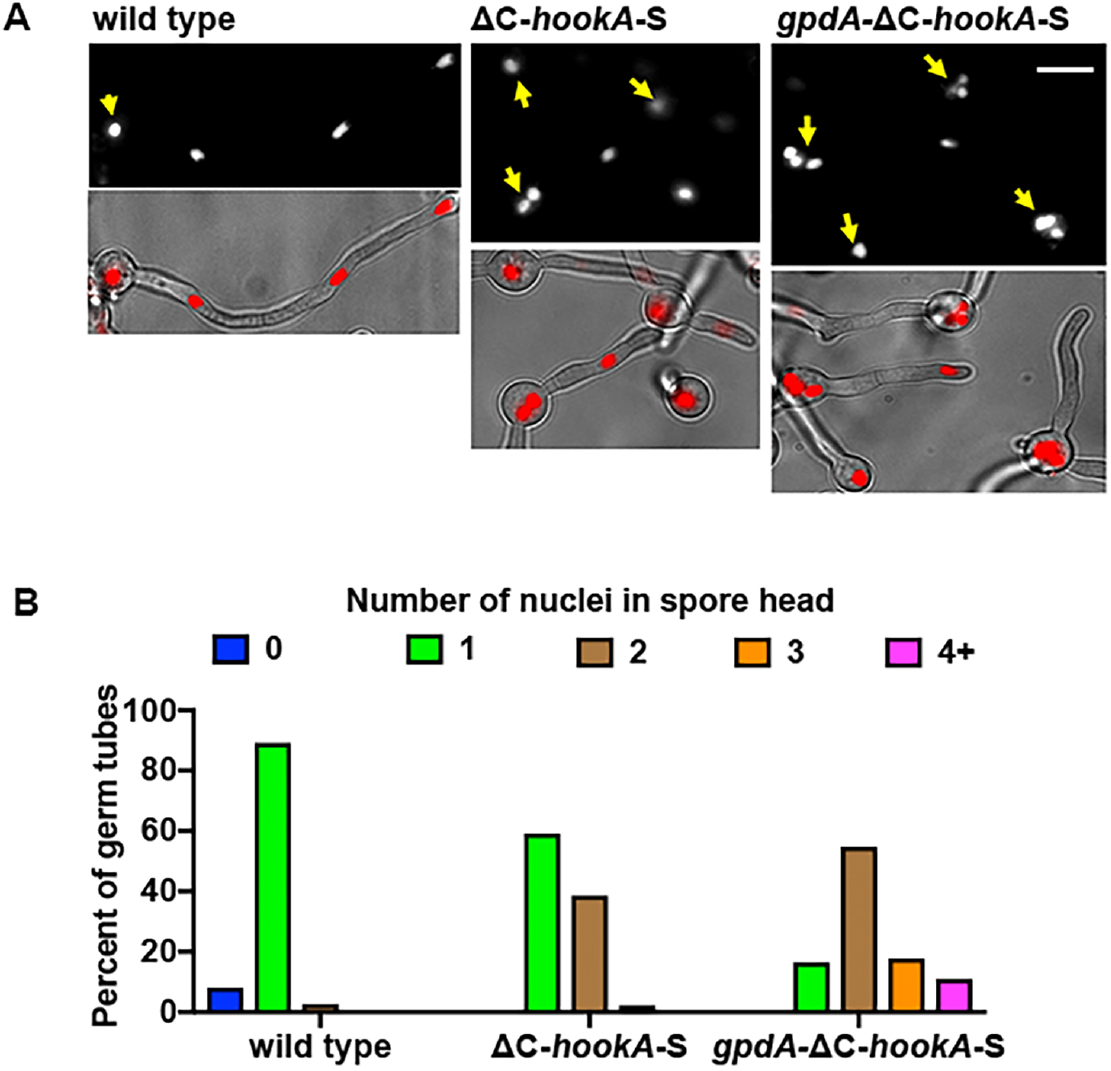
Overexpression of ΔC-HookA-S via the *gpdA* promoter causes a partial defect in nuclear distribution. (A) Nuclear distribution in the *gpdA*-ΔC-*hookA*-S strain in comparison to that in a wild-type control or a strain containing ΔC-*hookA*-S under the control of *hookA*-s endogenous promoter. Cells were grown in minimal glycerol medium at 32°C for overnight. Nuclei are labeled with NLS-DsRed. Bright-field images merged with the red nuclei are shown below. Yellow arrow indicates the position of spore head from where the germ tube emerges. Bar, 5 μm. (B) A quantitative analysis on the percentage of germ tubes containing 0, 1, 2, 3 or ≥4 nuclei in the spore head (n=74 for wild type, n=88 for the ΔC-*hookA*-S strain and n=73 for the *gpdA*-ΔC-*hookA*-S strain). The mean ranks of the three data sets are significantly different from each other (p<0.0001 for all comparisons) based on a nonparametric test that assumes no information about the distribution (unpaired, Kruskal-Wallis test, Prism 7). Note that the *gpdA*-ΔC-*hookA*-S strain exhibited an obvious but partial nud (nuclear distribution) phenotype, since a typical *nud* mutant like *nudA1* would have 4 or more nuclei in the spore head in 100% of the germ tubes.

**sFigure 2.**
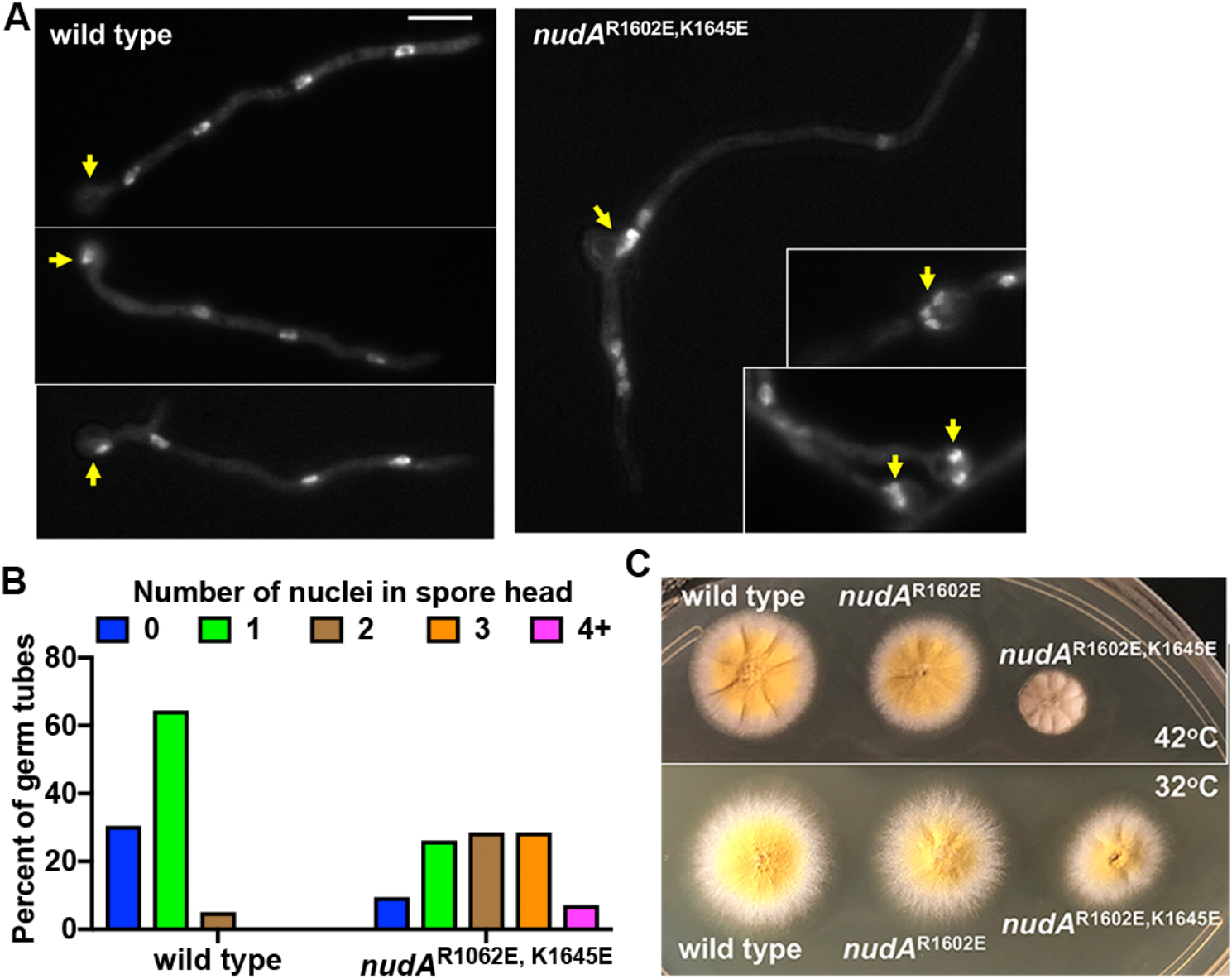
Phenotypic analyses of the phi-opening mutants. (A) The *nudA*^R1602E, K1645E^ mutant exhibited a partial nuclear-distribution defect. Cells were grown in minimal glucose medium at 37°C for eight hours. Nuclei were stained with DAPI. Yellow arrow indicates the position of spore head from where the germ tube emerges. Bar, 5 μm. (B) A quantitative analysis on the percentage of germ tubes containing 0, 1, 2, 3 or ≥4 nuclei in the spore head (n=59 for wild type and n=42 for the *nudA*^R1602E, K1645E^ mutant). The mean ranks of these two sets of data are significantly different (p<0.0001, two-tailed) based on a nonparametric test that assumes no information about the distribution (unpaired, Mann-Whitney test, Prism 7). Note that the *nudA*^R1602E, K1645E^ mutant exhibited a partial nud (nuclear distribution) phenotype, since a typical *nud* mutant like *nudA1* would have 4 or more nuclei in the spore head in 100% of the germ tubes. (C) Colony phenotypes of the *nudA*^R1602E^ and *nudA*^R1602E, K1645E^ mutants at 32°C and 42°C. The strains were grown for 2 days. Note that the *nudA*^R1602E, K1645E^ mutant is slightly temperature-sensitive because its defect in colony growth is more obvious at 42°C than at 32°C.

**sFigure 3.**
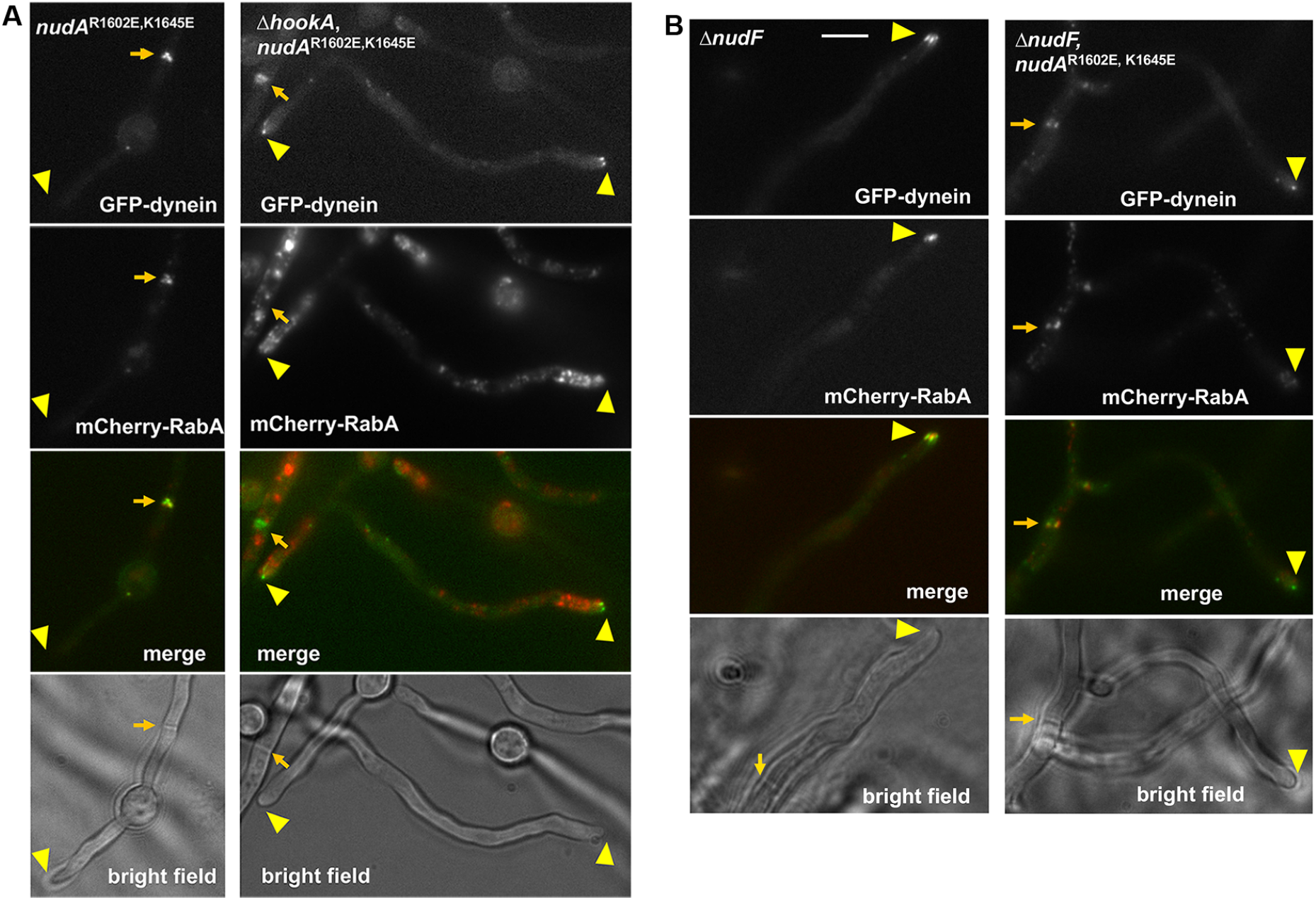
Early endosomes and dynein with the phi-opening mutations in the Δ*hookA* and Δ*nudF* mutants. (A) Images of GFP-dynein and mCherry-RabA-labelled early endosomes in the *nudA*^R1602E, K1645E^ phi-opening mutant and the Δ*hookA*, *nudA*^R1602E, K1645E^ mutant. Hyphal tip is indicated by a yellow arrowhead and septum by a light brown arrow. Note that GFP-dynein with *nudA*^R1602E, K1645E^ accumulates at septa with mCherry-RabA-labelled early endosomes. However, in the absence of HookA (Δ*hookA*), GFP-dynein with *nudA*^R1602E, K1645E^ accumulates at MT plus ends as comets as well as septa, but mCherry-RabA signals do not co-localize with those of GFP-dynein. (B) Images of GFP-dynein and mCherry-RabA-labelled early endosomes in the Δ*nudF* mutant and in Δ*nudF* with the phi-opening mutations *nudA*^R1602E, K1645E^. Hyphal tip is indicated by a yellow arrowhead and septum by a light brown arrow. Note that GFP-dynein with *nudA*^R1602E, K1645E^ is able to accumulate at septa with mCherry-RabA-labelled early endosomes even in the absence of NudF/LIS1. In addition, although plus-end dynein signals are still present, early endosomes are no longer accumulated at the hyphal tip in the Δ*nudF* mutant. Bar, 5 μm.

**sFigure 4.**
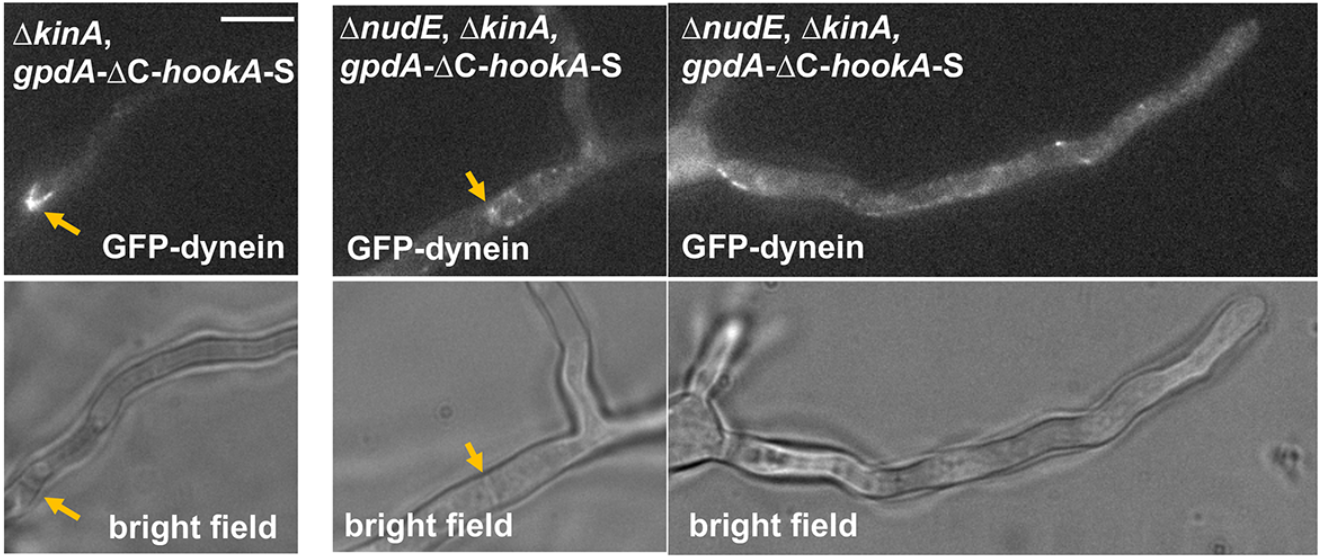
Images of GFP-dynein HC in a strain containing Δ*kinA* and *gpdA*-ΔC-*hookA*-S (left panels) and a strain containing Δ*nudE, ΔkinA* and *gpdA*-ΔC-*hookA*-S (right panels, showing two examples). Note that adding the Δ*nudE* mutation causes the septal accumulation of dynein to be less conspicuous and dynein along MTs to be more obvious. Septum is indicated by a light brown arrow. Bar, 5 μm.

